# Use of equine H3N8 hemagglutinin as a broadly protective influenza vaccine immunogen

**DOI:** 10.1101/2024.08.30.610504

**Authors:** David Verhoeven, Brett A. Sponseller, James E. Crowe, Sandhya Bangaru, Richard J. Webby, Brian Lee

## Abstract

An efficacious universal influenza vaccine remains a long-sought goal for human health as current vaccines suffer from shortfalls such as mid to low efficacy and the need for yearly revisions in strains use to account for viral drift/shift. Horses undergo bi-annual vaccines for H3N8 equine influenza virus, and surveillance of sera from vaccinees demonstrated very broad reactivity and neutralization to many human seasonal influenza strains. Subsequently, vaccination of mice using the equine H3N8 Kentucky/1/91 strain of equine H3N8 vaccine induced similar broadly reactive and neutralizing serum antibodies to human seasonal strains and to high pathogenicity avian influenza strains. Challenge of mice vaccinated with equine H3N8 or recombinant hemagglutinin (HA) based on the same strain protected vaccinees from high-dose lethal virus challenges. Protection was associated with the presence of neutralizing antibodies to the HA head, esterase, and stem regions that was similar to that of antibodies generated in horses after vaccination. Vaccinated ferrets exhibited similar broadly reactive serum antibody responses that protected vaccinees from clinical signs of infection and viral-induced histopathology after challenge with influenza A/07/2009 (H1N1) pandemic virus. Taken together, these data suggest that vaccination with equine H3N8 vaccine induces broad protection against influenza without the need for non-influenza viral vectors, multiple HAs, or foreign protein scaffolds common to other universal influenza vaccine candidates.

## Introduction

Seasonal influenza, and especially pandemic influenza viruses, continue to be one of the largest public health concerns of the 21st century. Seasonal influenza strains cause 3 to 5 million severe infections and 500,000 deaths yearly (WHO). Circulating influenza strains are very diverse because they are capable of undergoing both antigenic shift and drift, thus making vaccine design difficult. Occasional highly lethal zoonotic infections demonstrate that humans remain at constant threat of these viruses crossing the species divide and causing pandemics ^1^. Moreover, strains from subtypes with potential pandemic such as H5N1, H5N8, H7N2, H10N8, and H9H2 are in circulation in permissive species with the possibility for mutations or reassortment to allow crossing of the species barrier ^2^. The number of influenza strains that have crossed from birds into swine has increased dramatically in the last few years, further increasing the likelihood of crossing into humans ^3–6^. Thus, the discovery of a vaccine that can elicit protective immune responses against antigenically diverse strains is paramount for human health. Seasonal vaccination remains the best defense against influenza available, however, current vaccines have limited efficacy and provide a narrow breadth of protection ^7^.

Current licensed influenza virus human vaccines contain H1N1 (phylogenetic group 1 hemagglutinin), H3N2 (phylogenetic group 2 hemagglutinin), and 1-2 influenza B virus components. These vaccines are efficacious for closely matched strains predominantly by eliciting antibodies recognizing type-specific epitopes in the globular head domain of HA ^8^. Currently licensed vaccines are comprised of cold-adapted “attenuated” (LAIV), inactivated (IV), recombinant baculovirus-expressed HA, or virosome delivery of influenza peptides ^9,10^. IV still remains the most commonly administered vaccine, but efficacy is generally quite moderate ^11,12^, with LAIV often giving comparable protection. Recently, LAIV failed to protect from infection as demonstrated by the failure of the 2013-14 H1N1 pnd09 component of the vaccine (CDC) ^13^. As typified by the emergence of H1N1 pnd09 virus in the 2008-09 influenza season, some vaccines based on early season prediction strategies fail to prevent strains that emerge late in the influenza season. Furthermore, mutations in the predicted strains can further limit the protection afforded by current influenza vaccines as typified by the H3N2v2 that emerged in 2012-13 ^14^. Due to the limited efficacy of current influenza vaccine technology, predicting the next year’s relevant influenza strain or variance in antigenicity is not precise. Vaccine mismatches, antigenic variants and viral evolution of novel highly pathogenic strains such as avian H5N1, H7 and swine H1 and H3 variants complicate vaccine design. Vaccine failures due to antigenic mismatch and the pandemic threat from emergent viruses from reservoir hosts infected with avian or swine viruses warrant the development of influenza A vaccine candidates that induce broader protection. A universal influenza vaccine capable of providing broad protection against group 1 and group 2 influenza viruses would be ideal, but a broadly protective vaccine that extends coverage to most seasonal strains and the most likely pandemic strains is desirable.

Neutralizing antibodies (nAbs) generally bind to epitopes on the highly variable globular head domain of the HA, inhibiting receptor binding and mediating strain-specific hemagglutination inhibition (HAI) activity. Thus, nAbs to H1N1 pnd09 often only neutralize that same virus strain and are not broadly neutralizing Abs (bnAbs). Conversely, the stem domain of the HA (sHA2) ^15^ is relatively conserved but less immunogenic ^16^. sHA2 binding Abs can mediate HAI activity by inducing conformational changes within the head region. These mAbs have broadly neutralizing activity identified ^17,18^. Humans also recognize antigenic sites in the trimer interface domain that are broadly protective ^19–21^. These observations suggest that a vaccine that can induce such Abs would be more efficacious against divergent HA subtypes. However, vaccine approaches that seek to generate HA stem binding Abs may need to carefully ensure that the right Ab profiles are generated, as certain profiles of sHA2 binding Abs are associated with increases vaccine-enhanced respiratory disease in pigs ^22^. Whether or not enhanced disease can be induced in other species is uncertain. In addition, there is evidence that influenza may be able to modify its HA stem region to avoid stem-binding Abs ^23^, although this is unusual and occurs at a fitness cost for virus replication.

An optimal future universal vaccine may be a vaccine that targets conserved antigenic sites within both the head and the stem regions. A collection of nAbs that bind to a broad range of influenza virus strains and subtypes by binding to the sHA2 domain have been characterized ^24,25^, however a strategy of eliciting cross phylogenetic group HAI Abs via strain-specific vaccination has been difficult. Despite significant sHA2 conservation between similar groups of influenza strains, research to discover nAbs with activity against group 1 and group 2 viruses shows that B cells specifying these nAbs are rare. However, bnAbs can occur in humans that cross-react with H5N1 viruses ^26^, and generally are elicited by H1N1 pnd09 infections. Despite many common epitopes, infections with H3N2 viruses do not usually elicit broad protection against other antigenically divergent H3N2 strains, nor does H1N1 infection provide protection from similar subtypes, suggesting that conserved epitopes are not a major component of the typical immune response in healthy subjects with prior influenza exposure.

We recently discovered an alternative immunogen, based on equine H3N8 virus, which can elicit bnAbs by directing antibodies toward multiple HAs. We discovered that the equine HA3 antigen elicited HAI Abs toward many human group 1 and 2 viruses. We did not find equine H1N1 pnd09 infection, but did find it in cats, prompting further equine sera HAI and neuraminidase ELISA testing. Neutralization activity in equine serum with no evidence of infection by other strains prompted us to test vaccinations in mice and ferrets. This new immunogen elicited a favorable protective immune profile that targets multiple HA sites and cross-protects from heterosubtypic virus challenge in multiple species.

## Materials and Methods

### Challenge viruses

Challenge viruses were obtained from BEI Resources or the International Reagent Resource (IRR) as reassortantss with H1N1 A/PR/8/34 virus. BALB/c mice were inoculated with virus by large volume intranasal delivery to deliver virus to the lower respiratory tract, and lungs were harvested 3 days post-inoculation. Virus was passaged in mouse lung a total of 5 serial passages. The recovered strains then were expanded in 10-day old embryonated eggs (Charles River). Two additional H1N1 A/California/07/2009 (Cali/09) human isolates and an H3N2 A/Texas/50/12 isolate also were obtained from IRR and expanded similarly in eggs.

### Vaccine antigen

Live attenuated equine influenza virus (LAIV), based on the equine H3N8 Kentucky/1/91 strain, was obtained from Merck & Co. and expanded at 28 °C in 10-day old embryonated chicken eggs for 3 days. H1N1 A/California/07/2009 and H3N2 A/4230/2014 and H3N2 A/Texas/50/12 were passed serially in Madin-Darby canine kidney (MDCK) cell monolayer cultures at 37, 35, 30, and 28 °C before expansion in embryonated eggs, as described above for H3N8. These viruses were confirmed to grow in MDCKs at 28 °C to match the H3N8 LAIV vaccine. rHA was generated from H3N8 Kentucky/91 or H1N1 A/PR/8/34 viruses by overlapping synthesis of oligonucleotides (Genscript) that removed the transmembrane domain of the protein but inserted a flexible glycine/serine peptide spacer, followed by a T4 fold-on domain, and a 6× His tag. The sequence was subcloned into a baculovirus shuttle vector (Mirus Biotech) and used to produce infectious baculovirus according to the manufacturer’s instructions using the FlashBac Ultra system (Mirus Biotech) and transfection into Sf21 cells (Clontech).

After 5 days, the virus was titrated using a commercial titration kit (Clontech) by qPCR. HiFive cells (Invitrogen) then were inoculated with an MOI of 5 for two days before cleavage with N-tosyl-L-phenylalanine chloromethyl ketone (TPCK) treated trypsin. rHA was purified on Ni-NTA columns and dialyzed for 2 days with PBS using 4 buffer exchanges. rHA was routinely run on a native gel and verified to contain a ∼230 kDa protein that reacted with LAIV- immunized or PR8-immunized animal sera. Alum adjuvant was purchased from Invivogen and mixed with immunogen in PBS buffer 30 minutes prior to injection. Furthermore, vaccine stocks were tested for the presence of other viruses by H1 or H3 RT-PCR detection primers, and we only detected H3 amplicons or H1 amplicons in the matched vaccines (data not shown). In some experiments we directly compared H3 vaccines (*i.e*., H3N2 versus H3N8) to determine if responses were just from H3 immunizations. In others, we compared H3N8 to a mix of H1/H3 to simulate a human vaccine without the B strains.

### Animals

All procedures were approved by the Iowa State University Institutional Animal Care and Use Committee and Institutional Biosafety Committee. Euthanasia of mice and ferrets were consistent with the American Association of Veterinarians guidelines.

### Mice

Female BALB/c mice, 6 weeks of age, were purchased from the Jackson Laboratory and housed under ABSL2 conditions at Iowa State University. Mice were vaccinated twice with 25 μg of HA containing LIAV H3N8 or a mix of H1N1 A/07/2009 and California A/H3N2/4230/2014 or recombinant HA (rHA) by intramuscular injections containing alum.

Three weeks after the second vaccination, mice then were inoculated with influenza strains (2 to 4 × LD_50_ dose) H1N1, H3N2 in 40 µL of PBS under isofluorane gas. Mice were euthanized after >30% body weight loss or when exhibiting signs of respiratory distress.

### Horses

Horse sera from horses aged 2 to 10 years old were obtained retrospectively from the Iowa State University Diagnostic Laboratory. In addition, male and female horses, aged 25 weeks of age (receiving their first dose of FluAvert (Merck), were used from the Iowa State University herd prior to influenza vaccination. Horse peripheral blood samples were obtained by phlebotomy of the jugular vein 3 weeks post-H3N8 vaccine that was administered by the intranasal route according to the manufacturer’s instructions (Merck).

### Ferrets

Male ferrets, aged 4 weeks of age that were castrated and vaccinated for rabies and canine distemper virus, were purchased from Marshall and housed under ABSL2 conditions at Iowa State University. Ferrets were vaccinated thrice with 50 μL of H3N8, rH3 from H3N8, or H3N2 Hong Kong/2014 containing 25 μg HA and 0.2 μg of alum in PBS per intramuscular injection. Another group was vaccinated with alum alone in PBS. Ferrets then were inoculated intranasally with human influenza strain H1N1 A/California/7/2009 (10^7^ TCID_50_) in 1 mL of PBS under isofluorane gas.

### ELISA

Recombinant HAs were obtained from BEI Resources or the IRR. Antigen was added to PBS at 0.1 μg/mL and plated into 384-well Maxsorb ELISA plates (Thermo Fisher) overnight at 4 °C. Plates then were blocked with 3% nonfat dry milk (NDM) in PBS for 1 hr at 37 °C. Plates then were washed 4 times with PBS with 0.5% Tween-20 (Thermo Fisher). Sera were diluted in duplicates with two-fold dilutions down the short sides of the plates in NDM and incubated for 1 hr. Plates then were washed and anti-horse or anti-mouse IgM (Southern Biotech) or anti-horse (Southern Biotech) or anti-mouse IgG HRP (Biolegend) was added (1:10,000) in NDM and incubated for 1 hr at 37 °C. Plates were then washed and 3,3′,5,5′-tetramethylbenzidine (TMB) (Thermo Fisher) was added before reading optical densities. For the calculation of endpoint titers, a cutoff was determined as 3× the standard deviation of non-specific murine sera.

### Microneutralization assays

Murine, ferret, or equine sera were incubated 1:3 with receptor-destroying enzyme (Sigma) overnight at 37 °C followed by heat inactivation at 56 °C for 30 minutes. Influenza virus (100 TCID_50_/well) was incubated for 1 hr at 37 °C with sera diluted in DMEM (Thermo Fisher) medium with penicillin/streptomycin and 1% BSA. The virus was then added to plates containing MDCK cells (IRR) at 85% confluency and allowed to absorb for two hours at 37 °C followed by washing with DMEM. Cells were then incubated with culture media in the presence of TPK treated trypsin at 1μg/mL (Sigma) for 3 days. Viral neutralization titers were determined by incubation with 0.5% turkey blood (Lampire Biologicals) and calculated as reciprocal titers.

### Hemagglutination inhibition

HAI titers were determined similar to other studies ^27^ using serially diluted plasma with 8 HAU per 50 μL of H1N1 or H3N2 viruses in PBS containing 0.5% Bovine Serum Albumin (Sigma) in V bottom assay plates (Thermo Fisher). Virus/ antibody was incubated for 0.5 hrs at room temperature and then 0.5% turkey or chicken blood cells containing 0.5% BSA were added and plates read within 45 minutes. Data were calculated as the reciprocal titer above agglutination.

### Antibody competition assays

Competition with serum antibodies was performed by ELISA as described previously ^28^. Briefly, 96-well plates coated with 1µg/ml of H3 A/Texas/50/2012 HA were blocked with 5% nonfat dry milk, 2% goat serum, and 0.1% Tween-20 in PBS. The plates were washed with PBS containing 0.1% Tween-20 and incubated with serum diluted (1:10) with PBS, followed by wash and incubation with 5µg/ml of either mAb H3v-47 or mAb H3v-95 or mAb CR9114. Anti-human IgG alkaline phosphatase conjugate antibody (Meridian Life Science, W99008A) was used for detection with phosphatase substrate and the corresponding optical density values were measured at 405-nm wavelength on a BioTek plate reader.

### qPCR for viral burdens

50 mg tissue slices were extracted by Tissue RNA extraction kit (Qiagen) according to the manufacturer’s directions. 50 ng of RNA was then subjected to one-step qRT-PCR (NEB) with a universal M1 probe conjugated to FAM and Iowa Black quencher (IDT) for 40 cycles.

### Immunohistochemistry

5 μM sections were cut from paraffin blocks and embedded. Slides were washed through xylene and rehydrated in a series of ethanol washed from 100% to 0% before blocking with bovine FBS (5%) in PBS for 1 hour at room temperature. Slides were then stained overnight at 4 °C with mouse anti-influenza blend (Millipore) in PBS with 1% BSA at 1:100 dilution followed by three washes in PBS containing 0.05% Tween 20. Slides were then stained with anti-mouse FITC (BioLegend) at 1:100 in PBS with BSA at room temperature for 1 hr. Slides were washed again and stained further with anti-FITC Alexa488 (Jackson Immuno) in PBS with BSA at 1:100 for 1 hr at room temperature. Slides then were washed again and VECTASHIELD with DAPI (Vector) was added prior to imaging. Microscope gain and contrast were set using the sham vaccinees and uninfected ferrets and then all slides were imaged with identical settings. Slides were photographed in random spots on a fluorescent ZOE microscope (Bio-Rad) and images merged with no further manipulations except for automatic contrast done in PowerPoint (Microsoft).

## Results

### Vaccination with H3N8 induced cross-reactive antibodies to group 1 and 2 viruses

Horse sera with limited known prior vaccination histories were screened initially for serum antibody reactivity by ELISA across multiple rHAs and were found to contain broadly-reactive antibodies of both the IgM and IgG isotypes (Fig 1A and B). This reactivity pattern included both group 1 and 2 viruses and multiple potential pandemic strains. To assess for the effect of prior influenza exposure on the broad reactivity pattern observed, we also tested for antibodies directed toward N1 from H1N1 Cali/09 and did not detect reactivity (data not shown).

**Figure 1.**
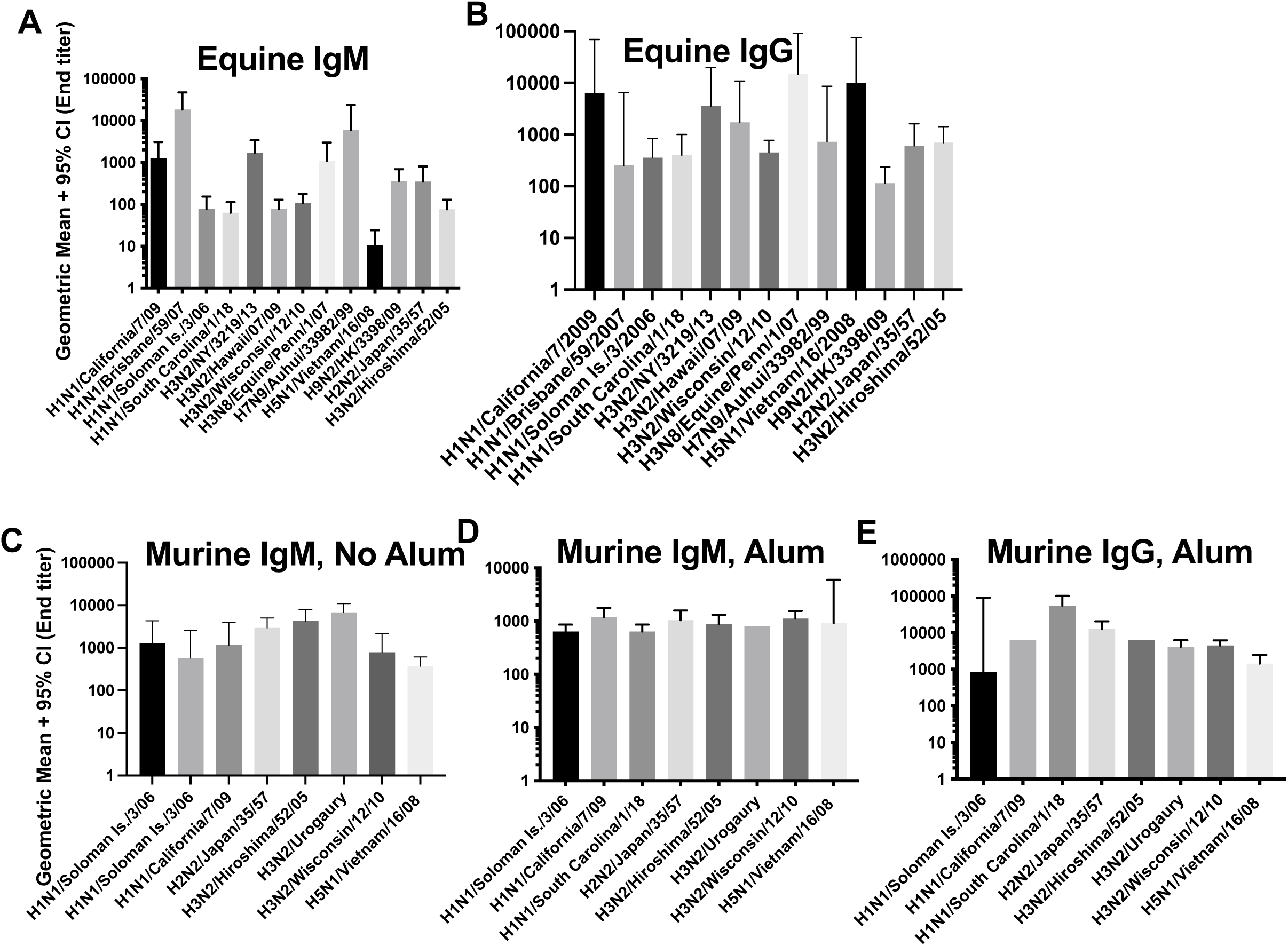
H3N8 vaccination leads to cross-reactive antibodies and heterosubtypic immunity. (A-B) Antibodies from horses were tested against a panel of rHAs (IRR Resources) from seasonal and avian flu strains by ELISA (n=15). **(C)** We vaccinated mice with two doses of our LAIV H3N8 without adjuvant and tested reactivity by ELISA again for IgM and IgG. **(D)** We then tested for IgM and IgG again after inclusion of alum in the immunization (C and D: n=5 for two reps each).

Next, we vaccinated mice with H3N8 live attenuated virus by intramuscular injection and found their serum antibodies exhibited a similarly cross-reactive pattern by ELISA after two vaccinations as that of the horses. We chose to inject the virus by IM injection to simulate a normal route for humans but also because we found equine H3N8 viruses would not infect mice at all doses tested. However, and in contrast with the horses, the murine reactivity was predominantly of the IgM isotype (Fig 1C) with little IgG, unlike the horses. We next tested whether including alum adjuvant in each intramuscular vaccination altered the isotype profile. We chose alum, knowing it’s a weaker adjuvant, since it is an approved and common adjuvant in humans. We found that alum altered the antibody profile toward a mixed IgM/IgG response (Fig 1D and 1E). All further mouse experiments used alum as an adjuvant in vaccine immunizations.

### Serum neutralizing antibody activity measured by HAI and microneutralization assays suggested immunogenicity of the HA head domain

HAI screening assays then were used with sera from vaccinated horses. These samples possessed broadly neutralizing activity toward multiple human seasonal influenza strains (Fig 2A). We then tested the responses of foals that had never been vaccinated with H3N8 and found similar neutralizing titers as the other horses with unknown vaccination histories (Fig 2B) but no responses prior to their first H3N8 vaccine.

**Figure 2.**
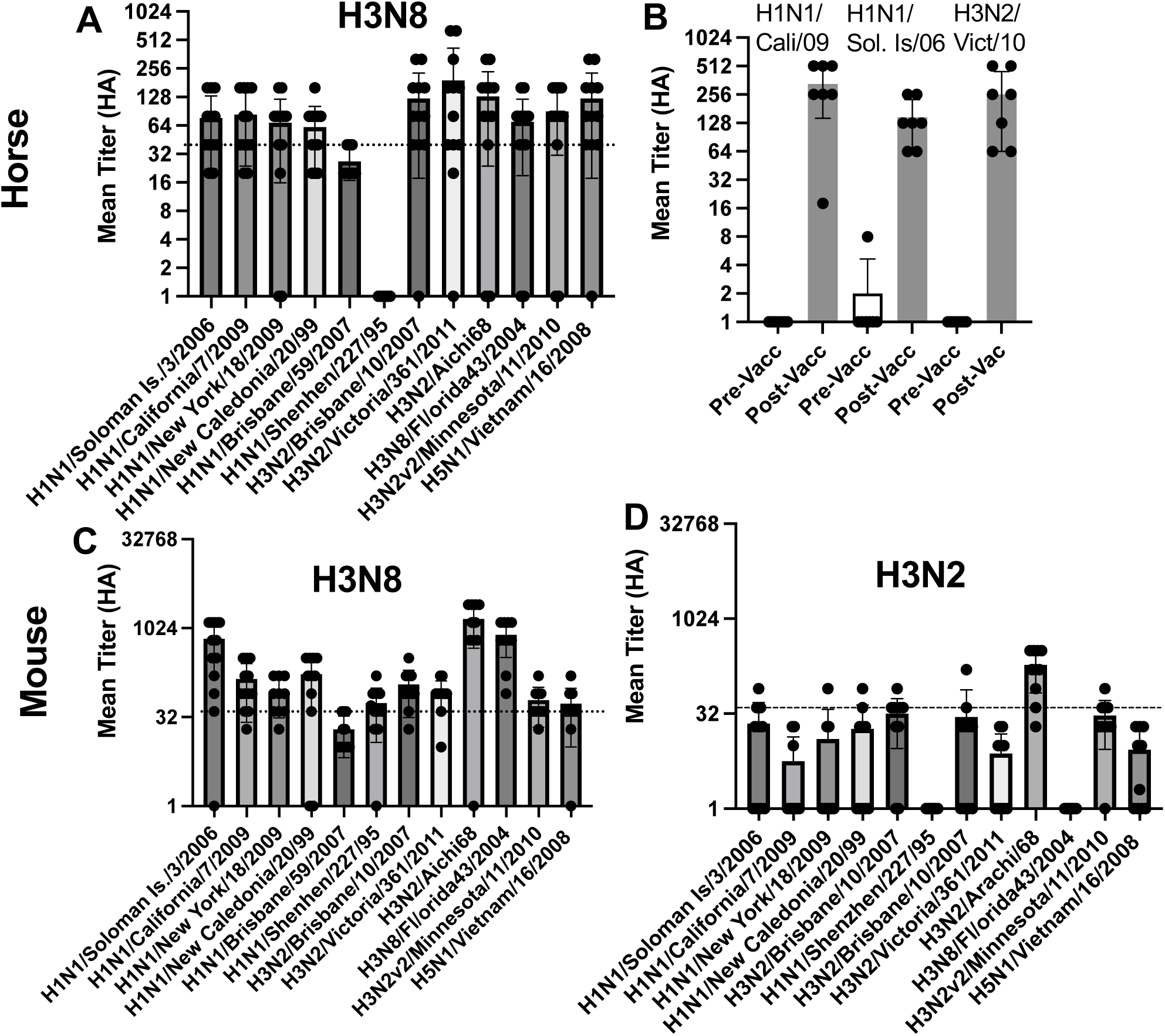
H3N8 induces bnAbs to human influenza strains. Sera from vaccinated horses were heat inactivated and RDE treated. **(A)** HAI titers were assessed across multiple strains of influenza (n=15). Line represents a protective reciprocal titer of 40. We do not show baseline pre-vaccination titers here since the horses are a mixed age population. **(B)** However, in a limited number of foals that we vaccinated for the first time, we did not find antibodies to any influenza antigen prior to vaccination and found equivalent HAI titers as their peers after vaccination (n=7, as finding vaccine-free horses is difficult). p<0.001 against unvaccinated foal sera. Mice received two doses of **(C)** H3N8 or **(D)** H3N2 (A/Texas/50/12) in alum, and HAI serum titers were tested across many strains of influenza. Line indicates a minimum protective reciprocal titer of 40. Similar influenza strains were at least p<0.001 between H3N8 and H3N2 vaccinees (n=5 for two reps).

In further agreement with the equine data, serum from vaccinated mice also had broad neutralizing activity across multiple human seasonal influenza strains (Fig 2C). Vaccination with a H3N2 virus (to match the H3 used in both vaccines) did not induce as broad of an HAI response as H3N8 vaccination, since the majority of responses were below an HAI titer of 40, with many having no response whatsoever (Fig. 2D). While we also determined that we could detect cross-reactive antibodies after one vaccine dose in mice, we could not detect cross-neutralizing antibodies until after the second vaccination (data not shown). We also tested sera from mice forneutralization activity and saw similar responses versus controls,thus validating our HAI results (Supplemental Figure 1)

### Mice and horses develop antibodies to the head, stem, and esterase regions after vaccination

We also compared the antibody reactivity of mouse vaccinee sera with that of multiple broadly neutralizing antibodies by ELISA and found a pattern similar to that of CR9114, although our antibody pools did not react to influenza B strains (Table 1). We then tested mouse vaccinee sera against the head (HA1) or stem (HA2) domains by ELISA and found that serum antibodies cross-reacted to both domains (Fig 3A). Finally, we used competition-binding assays to determine the approximate location of antibody binding against three HA antibodies H3v-95, H3v-47 and CR9114 specific to head, esterase-domain or stem, respectively^29^. We found that sera from horses or vaccinated mice could at least partially inhibit binding of antibodies directed to the head, esterase, or stem regions (Fig 3B). Mouse sera was not tested for head domain reactivity due to insufficient serum sample volume. However, the HAI data shown in prior figures clearly demonstrated mice also target the head region broadly. We also used the HAI and ELISA data against the various strains (Fig1/2) to examine for potential differences between neutralized and non-neutralized strains to map potential broadly neutralizing sites (Supplemental Fig 2). Future studies will be directed toward isolating such broadly reactive antibodies and characterizing them.

**Figure 3.**
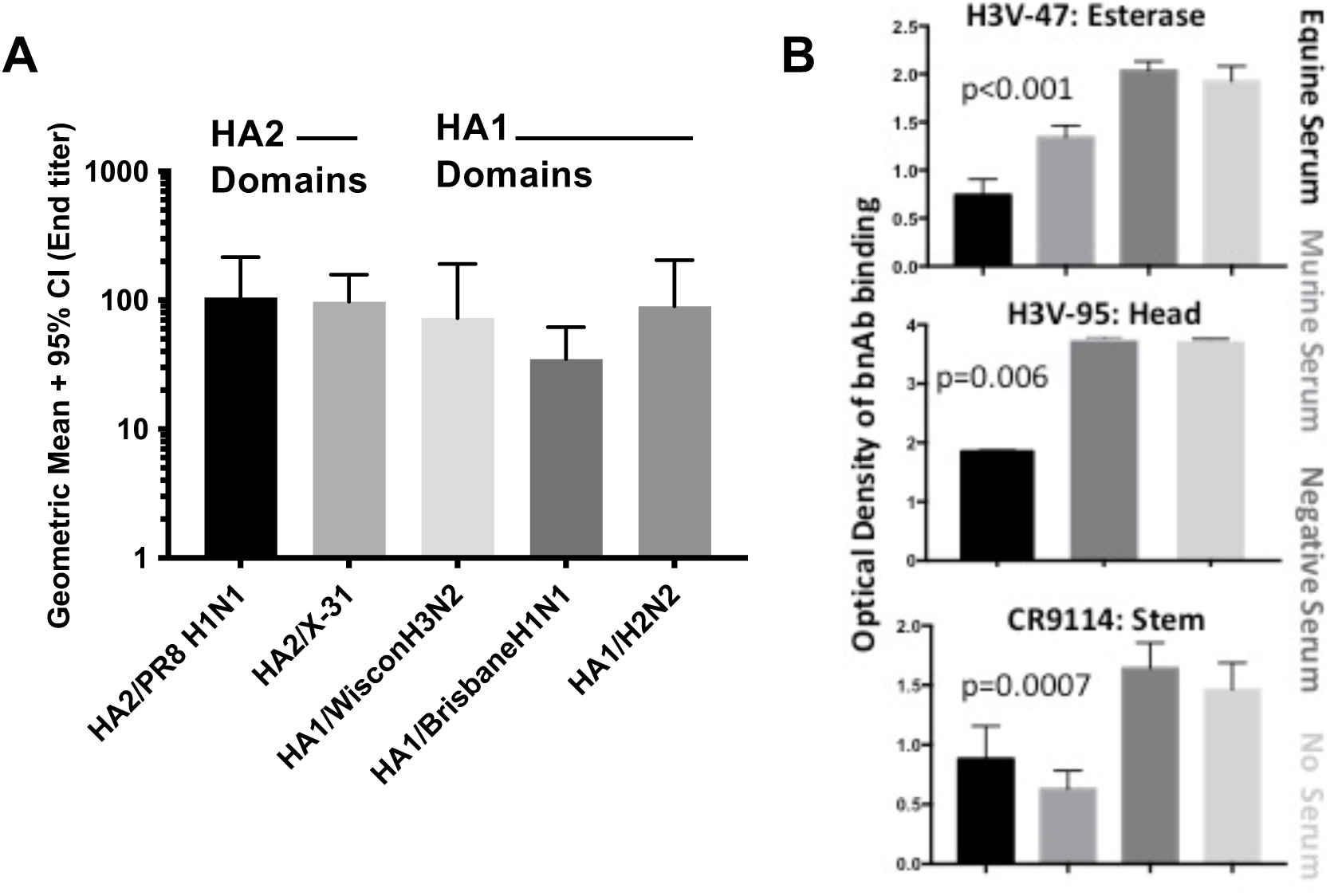
**Serum antibodies from vaccinated mice showed binding to HA stem and head domains**. **(A)** We performed ELISA for endpoint titers against headless HAs (sHA2) or recombinant HA1 portions of various rHAs (IRR Resources) (n=6 mice, 2 reps). HA-reactive antibodies were not detected in sham-vaccinated mice (not shown). p<0.05 or less for all comparisons to controls. **(B)** We performed ELISA competition-binding assays for antigen with pooled vaccinated mice (2x) or horse (1x) sera against panels of known bnAbs. Our sera competed for binding of antibodies specific for the esterase region (mAb H3v-47), head region (mAb H3v-95), or stem region (a recombinant mAb based on the sequence of CR9114) for binding to HA from H3N2 A/Texas/50/2012. Pools of n=5 mice or pools of n=10 horses were used.

**Table 1.**
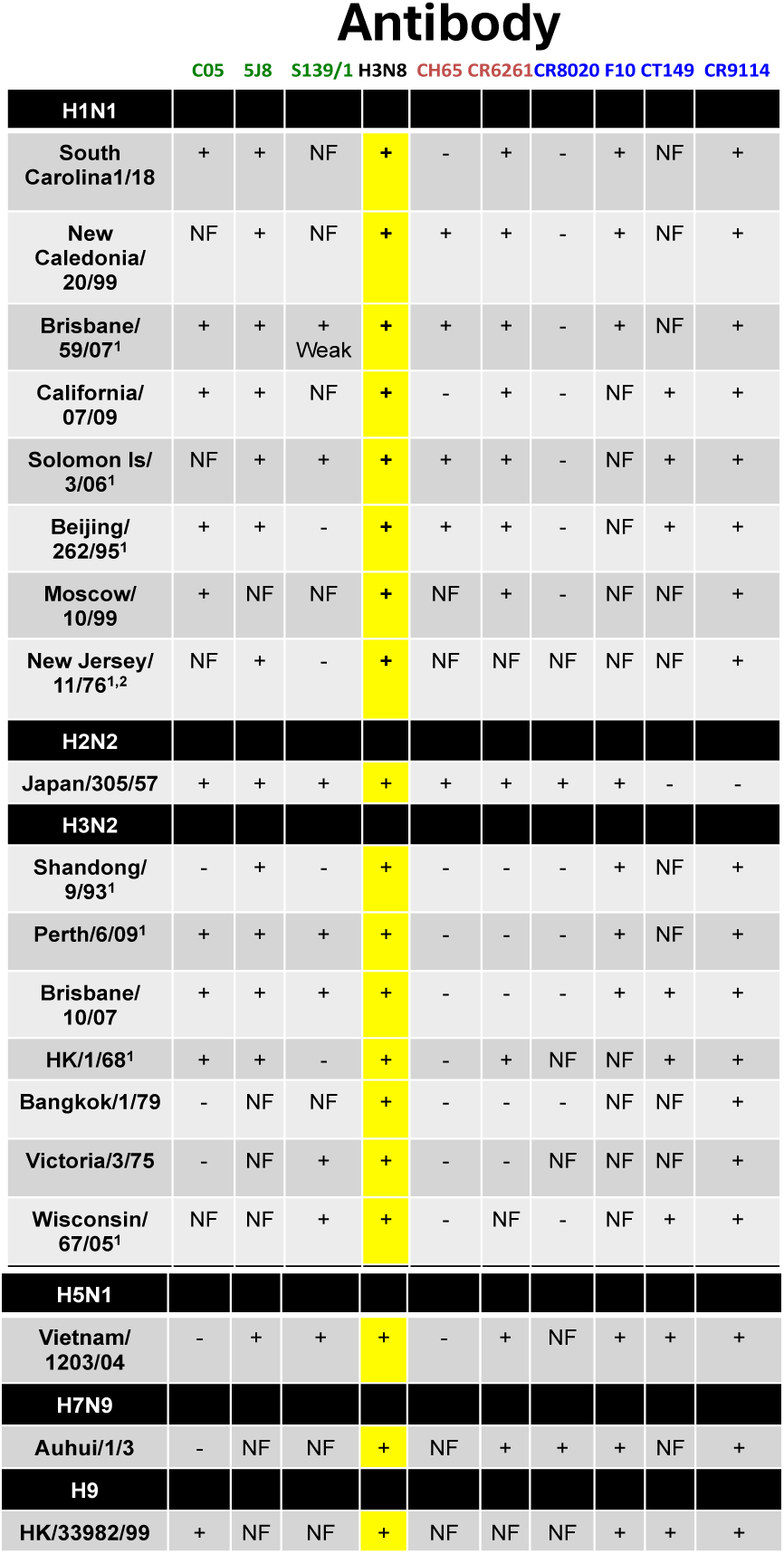
Cross-neutralization of H3N8. We compared neutralization results (>40 HAI) against the literature for head domain binding (green), esterase domain binding (red), or stem domain binding (blue) antibodies. H3N8 vaccination induced a polyclonal response, but nonetheless lead to higher cross-reactivity than that exhibited by bnAbs. Reactivity to type B strains was not detected in equine vaccinees.. NF= no evidence of neutralization found in the literature.

### Vaccination protected mice from heterosubtypic challenge

Mice were vaccinated either with attenuated H3N8, rHA, control viruses, or PBS alone, as above. To confirm that our rHA vaccine elicited similar HAI toward many seasonal strains like the whole H3N8, we first tested for neutralization after two vaccinations of either our H3N8 rHA or PR8 H1 rHAs (Fig 4A-B). In agreement with data in Fig 2C, we found neutralization of many seasonal strains for which PR8 rHA did not elicit neutralization (Fig 4B). We then challenged vaccinated mice with lethal doses of murine-adapted H1N1 or H3N2 viruses. We found that animals vaccinated with H3N8 or rHA were protected from lethality (Fig 4C-F). We next vaccinated mice similarly as before and used a 2×LD_50_ dose of A/PR/8/34 virus and found that H3N8- and rHA-vaccinated animals had what appeared to be less lung histopathology (Fig 5A-D) and had lower viral burdens in the lungs after challenge (Fig 5E).

**Figure 4.**
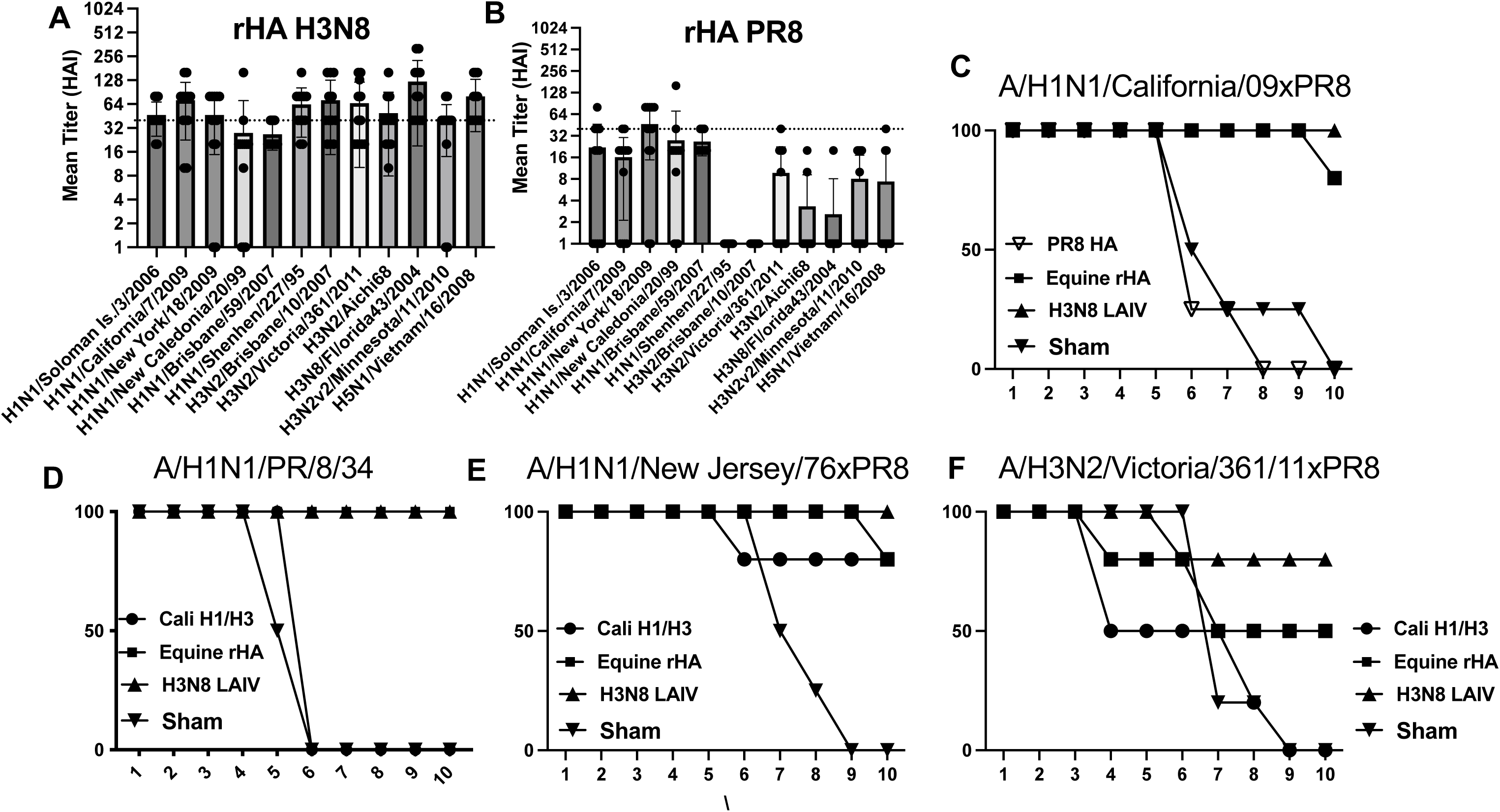
Protection against infection after vaccination. (A) We vaccinated mice as stated prior in results twice using rHAs from either H3N8 oand tested for HAI responses across many seasonal strains of influenza (n=5 for two reps). **(B)** Next, we tested for survival after H1N1 A/07/2009 challenge (4 × LD_50_). PR8 HA was used here instead of our H1/H3 controls since they contained H1N1 A/California/07/2009 virus (n=5 for two reps). **(C-F).** Next, we tested for survival after challenge with 2 × LD_50_ of viruses stated in the figures. Here we compared our H3N8 whole vaccine against our H3N8 rHA and a sham or control vaccine containing H1 and H3 viruses (n=5 for two reps for all animals). Per our animal protocol, we did need to use a 25% weight cut-off prior to euthanizing animals after challenge. p<0.01 for each of the survival curves between equine vaccinees (LAIV or rHA) and controls.

**Figure 5.**
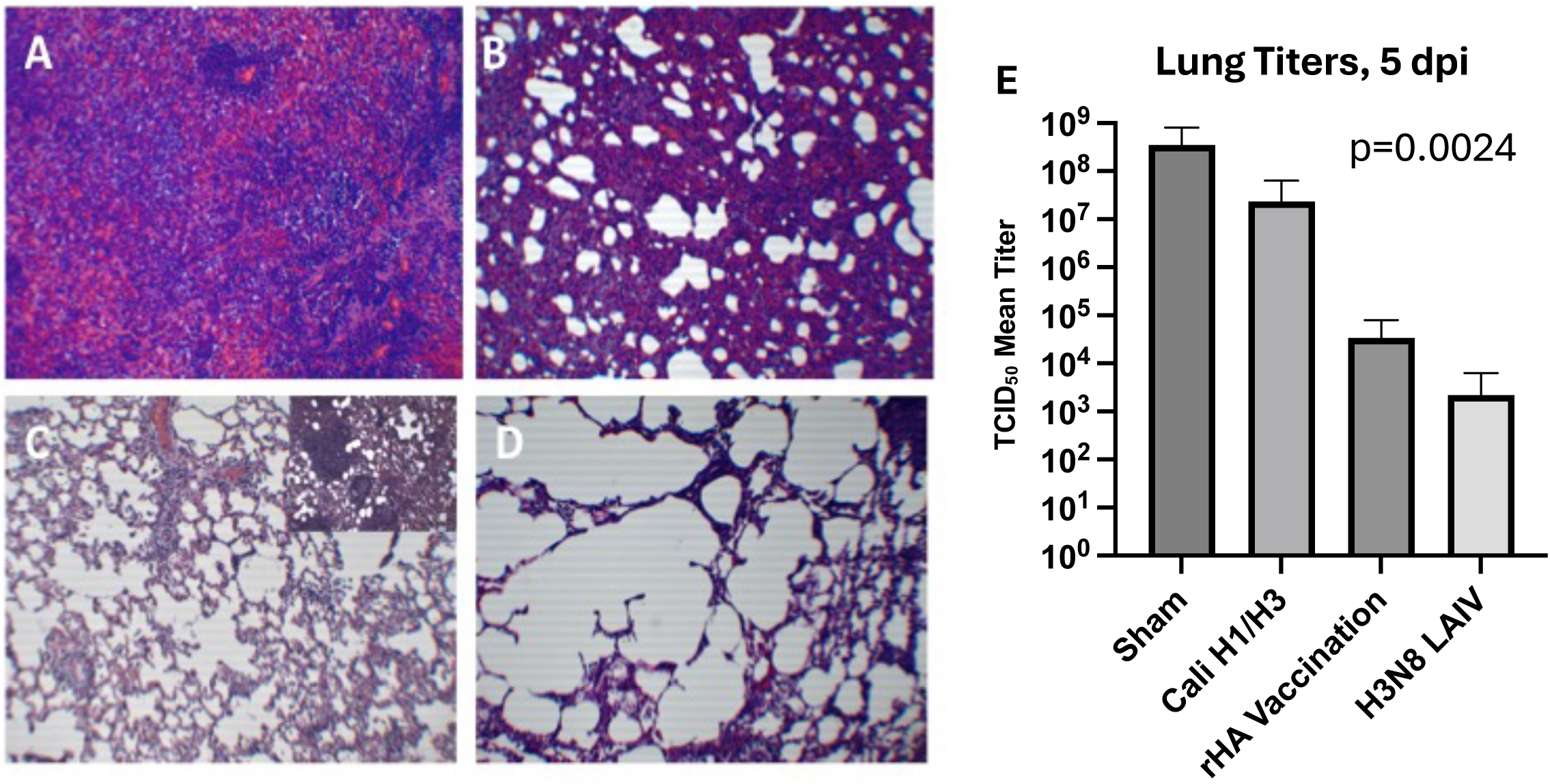
H3N8 protects from heterosubtypic challenge with lower viral loads. (A) We again vaccinated mice with either H3N8 whole virus, rHA form H3N8, sham, or our mix of H1/H3 Cali viruses. We assessed collected lungs after 6 DPI and stained with hematoxylin and eosin. (n=5 with two reps but representative photos are shown. Shown are (**A)** sham control, **(B)** H1/H3 Cali, **(C)** equine rHA, and **(D)** H3N8 LAIV. **(E)** Viral lung burdens were determined in 1mg of upper right lobe after 6 DPIs form the same mice as (A) by TCID_50_ assay.

### Immunogenicity and protective efficacy in ferrets suggest H3N8 protects from challenge in another species

Next, ferrets were vaccinated with rHA, LAIV, H3N2 A/Texas/50/12, or PBS sham by intramuscular vaccination as done in mice. HAI titers then were examined after one, two or three vaccinations and found to have increasing levels of HAI to H3N8 virus (Fig 6A) and cross-reactive HAI against H1N1 A/California/07/2009 after each vaccination (Fig 6B). We then challenged the ferrets four weeks after the third vaccination with 10^7^ TCID_50_ of H1N1 A/California/07/09 virus. Ferrets vaccinated with sham or H3N2 A/Texas/50/12 had significant lung consolidation similar to mice from Fig 5 (not shown). Ferrets vaccinated with rHA from H3N8 had mixed histopathology at 6 DPI, with most exhibiting areas of clearance but some areas with lung infiltration and consolidation that still appeared better than that in controls like Fig 5. Ferrets vaccinated with LAIV H3N8 had lung histology looked very similar to uninfected controls. Sham and A/Texas/50/12 inoculated ferrets also had higher sneezing and lethargy scores than equine vaccine recipients. We next stained those same lungs with pan anti-influenza A antibody (Millipore). We found that sham-immunized ferrets had wide-spread distribution of virus in the lungs (Fig 7A) while H3N2 A/Texas/50/12 immunized animals had less virus but still substantial virus antigen staining. In contrast, rH-immunized ferrets had significantly less but still detectable virus by staining, while it was more difficult to to find areas of viral staining in LAIV-vaccinated ferrets.. Viral burdens were determined by qRT-PCR on nasal samples, and we found that H3N8-vaccinated ferrets had the least viral burdens followed by rHA-vaccinated animals and then the controls in nasal swabs at 3 DPI (Fig 7B). Alternatively, viral titers were determined as TCID_50_ of 1 mg sections of upper right lobe lungs at 6 DPI (Fig 7C).

**Figure 6.**
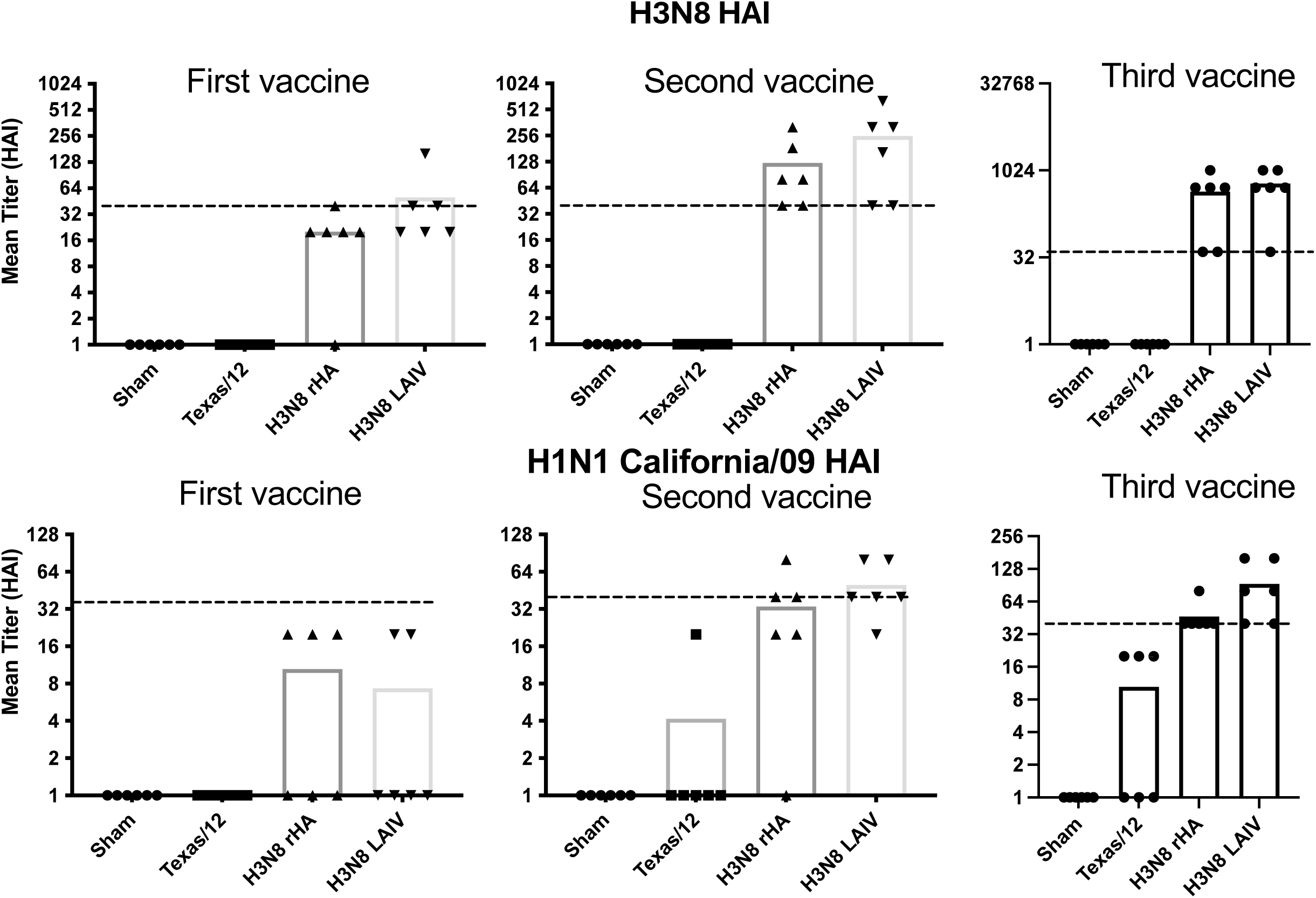
Vaccination in ferrets leads to cross-reactive HAI antibodies similar to that in other animal vaccinees. Ferrets received 3 doses of immunogens with alum and were then challenged 4 wks after the third vaccination. **(A)** HAI responses to H3N8 virus after each vaccination. p<0.001 between equine and control groups. **(B)** HAI responses to H1N1 A/California/07/2009 strain after each vaccination. Notice the third vaccine axis are different from subsequent vaccinations. p<0.001 between equine and control groups. Lines represent HAI titer of 40.

**Figure 7.**
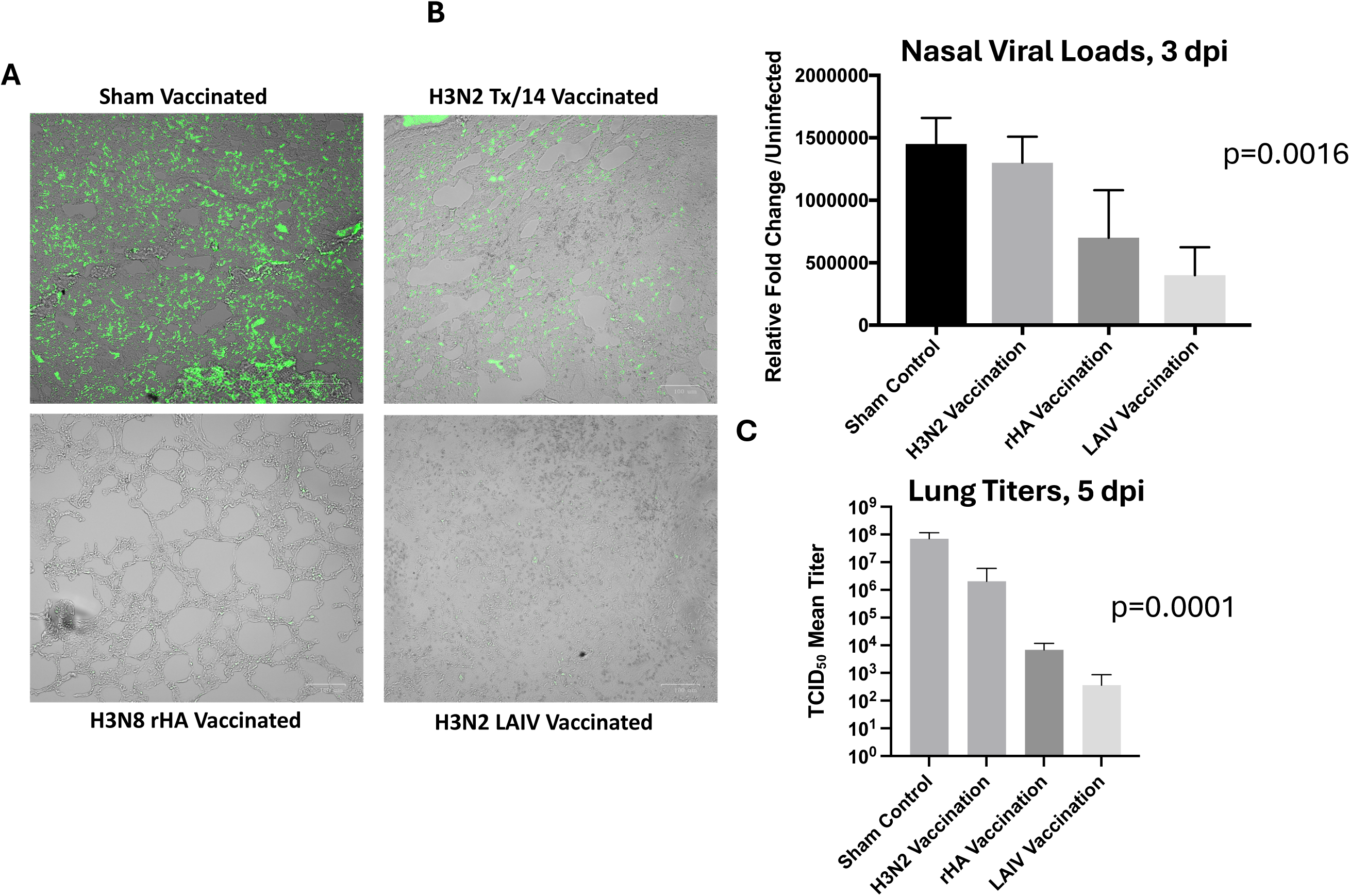
Reduced viral shedding in H3N8 vaccinees after challenge with H1N1 A/California/07/09. Ferrets rested three weeks post their third vaccine were then challenged with 10^7^ TCID_50_ of H1N1/Cali/09. **(A)** Ferrets were euthanized after 6 days and lungs stained with anti- influenza A antibody. A representative stain is shown for all groups. Brightfield images were merged with FITC images by the camera software (4× magnification). Green indicates the presence of influenza virus. **(B)** We assessed the viral burdens in nasal swabs at 3 days post-infection from the same ferrets using one-step qRT-PCR for a flu conserved M2 region. **(C)** 1 mg of lung tissue was homogenized and filtered, and then virus titer was determined by TCID_50_ assay.

## Discussion

There is an urgent need for a broadly protective influenza vaccine given the yearly circulation of strains that infect humans, the constant antigenic drift and possibility for an antigenic shift, and the threat of potentially highly pathogenic strains crossing into humans. In the present study, we demonstrated the efficacy of an equine influenza H3N8 virus to protect from a heterosubtypic influenza virus challenge in multiple animal species, suggesting further development of this novel antigen as a vaccine candidate is warranted.

Interestingly, dogs infected by H3N8 virus from cross-species exposure to infected horses were susceptible to infection by H3N2 in subsequent cross-species transfers, but only H3N8 viruses infect horses ^30^. While experimental infections with high doses of other viruses are possible in horses, those infections are generally attenuated as compared to H3N8 ^31^. While these observations could be due to differences in viral tropism, another possibility is that H3N8 exposure in horses makes them immune to other influenza viruses. Further evidence for this possibility is that prior to H3N8 adaption into horses, H7N7 viruses were the predominant infecting viral species, but that viral strain disappeared from horses after H3N8 crossed into horse populations (Murcia et al., 2011).

It has been previously reported that mice or ferrets vaccinated with a cold-adapted virus incorporating equine HA induced protection from equine influenza challenge ^32^, which shows suggested vaccination might prevent that particular zoonosis due to equine influenza virus. Here, we extend those findings to show that equine H3N8 antigens or epitopes might provide a promising component for a more broadly protective vaccine against human influenza. The previous studies used equine H3N8 Georgia/81 virus, and there are key differences with the influenza strain used here H3N8 Kentucky/1/91, as shown in Supplemental Figure 3.

Specifically, there are key amino acid substitutions in areas that we predict to induce bnAbs (*e.g.*, site S70P has no glycosylation site in the Georgia/81 vaccine, meaning a key immunodominant and non-protective epitope is exposed that might decoy the immune response near a site we hypothesize contributes to protection). As we continue to examine this vaccine candidate, testing other equine influenza viruses against heterosubtypic protection could be an interesting avenue of exploration.

The need for high concentrations of alum adjuvant to push the antibody profile from IgM- towards IgG-dominant suggests that a lack of CD4 T cell help may occur with this antigen, at least in mice. To help with this shift, we chose to study alum here since it is an approved adjuvant in human vaccines. Likely, other adjuvants might increase the level of HAI titers or induce a better IgG-dominance than what we observed with alum. In our preliminary studies, inclusion of cyclic dinucleotides adjuvants can increase the HAI titers observed at least 4-fold in mice (not shown), but we do not yet know about the effect of such adjuvants on the switch from IgM to IgG. Our goal in inducing class switching is to induce an antibody response of higher affinity and longer duration. IgM antibodies are known to be produced in the lungs after the challenge of mice and can have significant reactivity toward influenza, but their duration may be short-lived ^33^. In humans, neutralizing IgM antibodies may exhibit a longer half-life after influenza exposure, and thus IgM might contribute to clinical protection in humans ^34^.

The difference in protection between our baculovirus-expressed HA and the live attenuated virus likely is due to two factors: (1) the whole virus contains more antigenic proteins such as nucleoprotein (NP) and neuraminidase (NA) and (2) baculovirus-expressed HA protein generally needs much higher concentrations in the injection to confer similar efficacy as split vaccines. We saw at least some serum antibody cross-reactivity in NA-based ELISAs in our whole virus H3N8 vaccinees that we did not see in the serum of H3N2 vaccinees. Work to qualify and quantify any contributions from antibodies toward NA is of interest (Supplemental Figure 4, not an exhaustive testing across strains yet). We also did not examine cell-mediated responses toward the vaccine, and likely CD8 T cells directed toward the NP contribute to heterosubtypic protection observed in animals. Moreover, there is evidence in humans that CD8 T cells directed toward NP may impart some protection.

The need to use three doses of vaccine in ferrets stems from the fact that we did not see robust HAI titers toward H1N1 A/California/07/09 after two doses. This limited response could stem from a lack of prior exposure to influenza vaccines. Some H3N2 viruses elicit low immunity by intramuscular injection after the first dose versus intraperitoneal routes in mice. Alternatively, we may not have included a high enough dose to optimally boost the HAI titers after the prime and boosting doses, as we have evidence in our murine vaccinations that a third and fourth vaccination increased the total sera concentration of H3N8 reactive antibodies. This observation suggests that maximal immunogenicity is not achieved after two doses with the current regimen and formulation. In fact, we found in subsequent experiments that further boosting mice with another 2 doses of H3N8 or rHA increased antibody titers suggesting that stronger adjuvants or higher doses might increase the level of response. We also believe a similar trend might occur in our vaccinated ferrets, as likely even three doses of immunization did not maximize the full level of HAI we could have obtained.

Here, we demonstrated that equine H3N8 can induce broadly reactive and neutralizing antibodies to multiple human season influenza strains. Follow-on studies will compare rHA in higher concentrations or with other viral proteins (*i.e*., NP or NA) to determine whether we can bolster the protection afforded similar to that induced by the whole virus. We also are evaluating other antigen delivery methods including prolonged polyanhydride nanoparticle release. At present, we do not yet know if prior exposure to other influenza strains for vaccines changes the immunity imparted by this vaccine candidate. We also do not yet know how long the induced immunity lasts, and thus do not know whether an annual booster would be necessary to maintain immunity over time. We are exploring the antigenic sites recognized by the induced neutralizing antibodies and are actively generating hybridomas from vaccinees (equine and murine). While H3N8 virus is not very infectious for humans, the inclusion of the vaccine as a killed or rHA vaccine could foster protective responses. We demonstrated that the H3N8 virus or its HA is capable of affording broad protection for influenza A strains across multiple animal species and multiple seasonal influenza strains and consider this vaccine candidate worthy of further development.

Future studies will explore the use of other equine strains or HAs and testing higher doses or alternative adjuvants with the aim to bolster to enhance immunogenicity. Of course, cellular immunity form the vaccine might also contribute to protection and will need to be studied.

However, these studies establish proof of principle for the potential of this immunogen to induce broadly protective responses.

## Acknowledgments

The work was supported in part by a research grant from the Investigator-Initiated Studies Program of Merck Sharp & Dohme LLC. The opinions expressed in this paper are those of the authors and do not necessarily represent those of Merck Sharp & Dohme LLC. We further acknowledge the NIH Biodefense and Emerging Infections Research Resources Repository NIAID, NIH and Influenza Reagent Resource, Influenza Division, WHO Collaborating Center for Surveillance, Centers for Disease Control and Prevention for providing rHAs and viruses used in this study.

## Conflict of Interest

The vaccine used in this study is based on a veterinary vaccine (FluAvert) manufactured by Merck Sharp & Dohme LLC. However, Merck was not involved in the planning or execution of this study. DV and BS have a patent granted for the use of A/equine/Kentucky/91 for use as a universal influenza vaccine. J.E.C. has served as a consultant for Luna Labs USA, Merck Sharp & Dohme Corporation, Emergent Biosolutions, a former member of the Scientific Advisory Boards of Gigagen (Grifols), of Meissa Vaccines, and BTG International, is founder of IDBiologics and receives royalties from UpToDate. The laboratory of J.E.C. received unrelated sponsored research agreements from AstraZeneca, Takeda Vaccines, and IDBiologics during the conduct of the study.

---

**Supplemental Figure 1.**
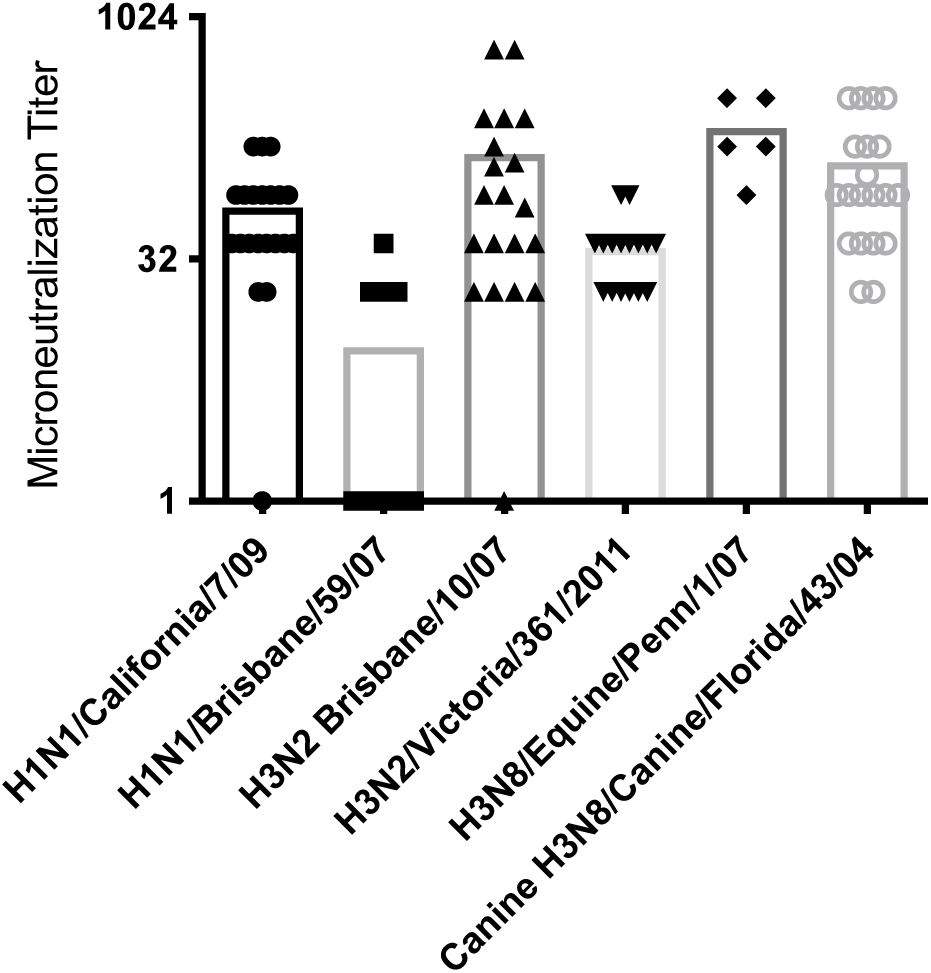
Microneutralization data confirms HAI data. We tested for microneutralization across multiple strains of influenza. (n= 7 to9 for two reps each).

**Supplemental Figure 2.**
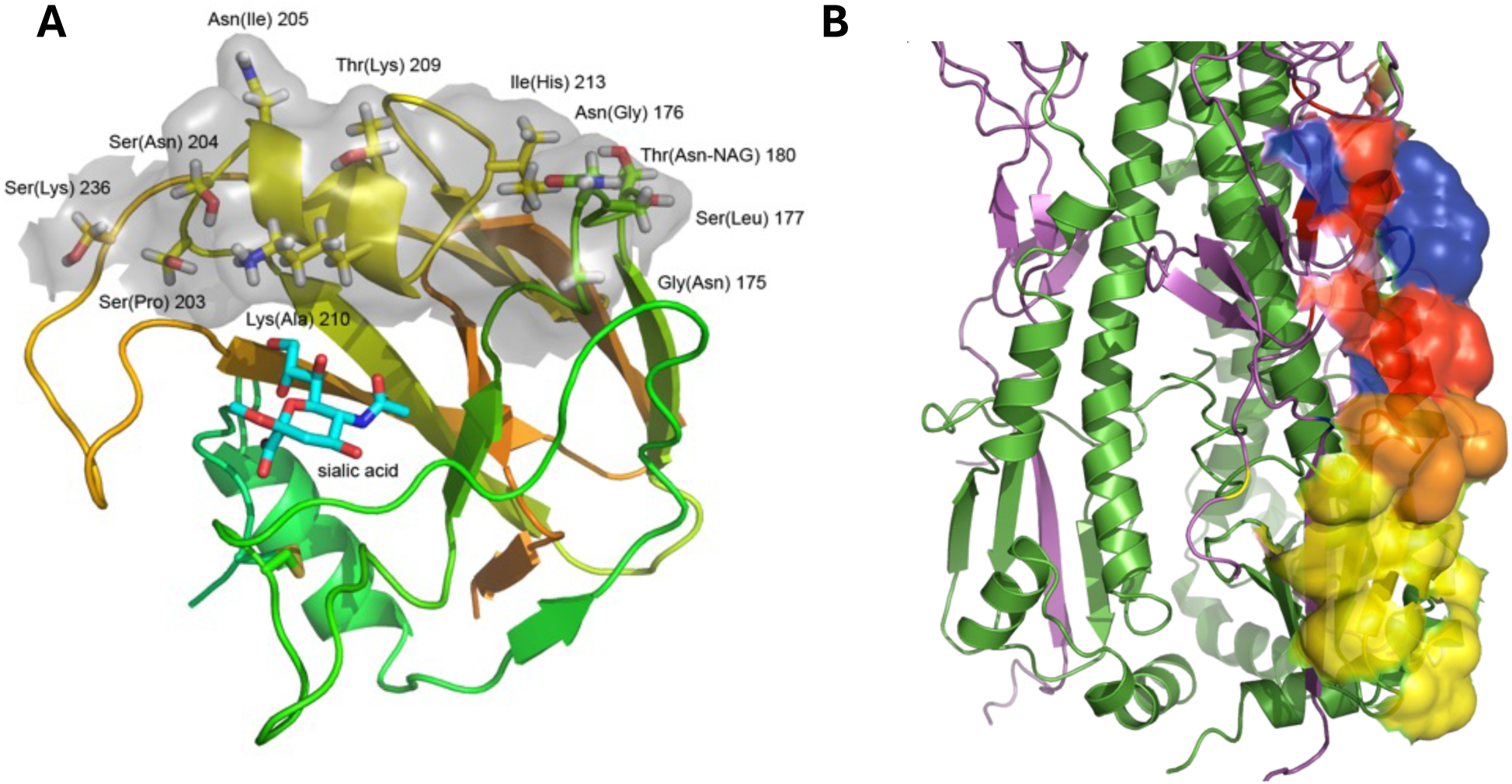
P**o**tential **conformational epitope from equine H3N8 HA. (A)** Sequence homology to the reactive human H1N1 (strain A/California/04/2009) and divergence from non-reactive human H1N1 (Brisbane, equine sera) was used to map the potential epitope to the HA1 head region. The epitope surface (grey shading) was mapped onto a homology model of equine H3N8 HA (strain A/Xuzhou/01/2013) with amino acids numbered according to the full-length HA sequence and H1N1 HA variants indicated in parenthesis. In addition to the sequence variants, a predicted glycosylation site at Asn180 may impair binding to this epitope. Antibody binding is expected to inhibit sialic acid binding. The location of the bound sialic acid is based on superposition with a known structure of HA (PDB ID: 1HGE) ^35^. The homology model of equine HA was calculated with RosettaCM using structural templates of HA from equine, canine and harbor seal H3N8 influenza viruses (PDB IDs: 4UO0, 4UO4 and 4WA1) ^36–38^. **(B)** There are overlapping stem epitopes for bnAbs C179 and CR6261, which bind to Group 1 HAs. There is considerable conservation between all HA strains within the HA2 portion of these epitopes, but not within the HA1 portion (particularly with the glycosylation site at Asn53 in equine/Kentucky/3/91/H3N8). Examination of the structure suggests that there is the possibility to recognize a cross-protomer epitope that includes the conserved HA2 and a neighboring HA1 beta hairpin across the trimer boundary. Examination of sequence shows that there is some conservation, but there is a glycosylation site in H1N1 strains at Asn40 that would block this site. The recognition site also could be similar to that of bnAb FI6v3, which has a similar epitope as that of CR6261, but adapts to the glycosylation site at Asn53 in H3 strains and minimizes interactions with the HA1 parts of the epitope (in particular there is almost no contact with the Asp306-Lys307-Pro308 loop, numbering from equine Kentucky H3N8). Another broad stem antibody CR9114 appears to bind similarly to FI6v3 and accommodates the Asn53 glycosylation site. So, if the binding we see is not due to multiple binding sites from more than 1 bnAbs, then an epitope that overlaps with that of CR6261 is possible, and the antibodies induced could adapt to strain variations and glycosylation. **purple** – HA head, **green** - sHA2, **red** - overlapping epitope surface of C179 and CR6261, **pink** - C179 only, **blue** - CR6261 only, **yellow** - CR8020 only, **orange** - overlap between all three antibody epitopes. Fl6v3 ^39^ is exceptionally broadly binding as are the preliminary screens of the H3N8 antibodies (see Table 1) under study.

**Supplemental Figure 3.**
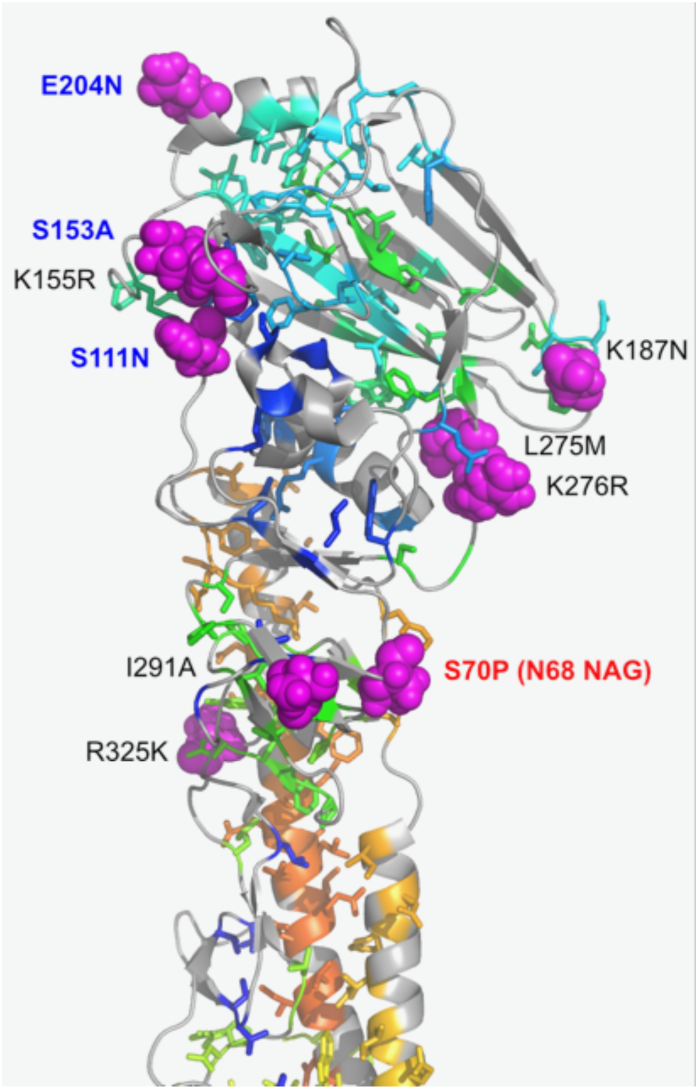
Divergent differences in equine HA antigen. While equine HA has 95 to 98% amino acid homology between all strains, there are some key differences that might make one more of an ideal immunogen. Here, we compared the sequence of equine Georgia/81 HA (shown) with that of our immunogen (Kentucky/1/91) and found key differences in amino acid substitutions. The S70P site indicated in red shows a difference in glycosylation at N65 (present in our immunogen). Magenta colored residues shows other key differences with our immunogen (labeling with our immunogen first followed by Georgia). These regions likely contain protective epitopes. The blue label color indicates areas of difference with other equine influenzas from the Florida lineage (*e.g.*, Sydney/6085/07). Thus, there are key sequence differences in equine HA proteins that may influence how the host responds.

**Supplemental Figure 4.**
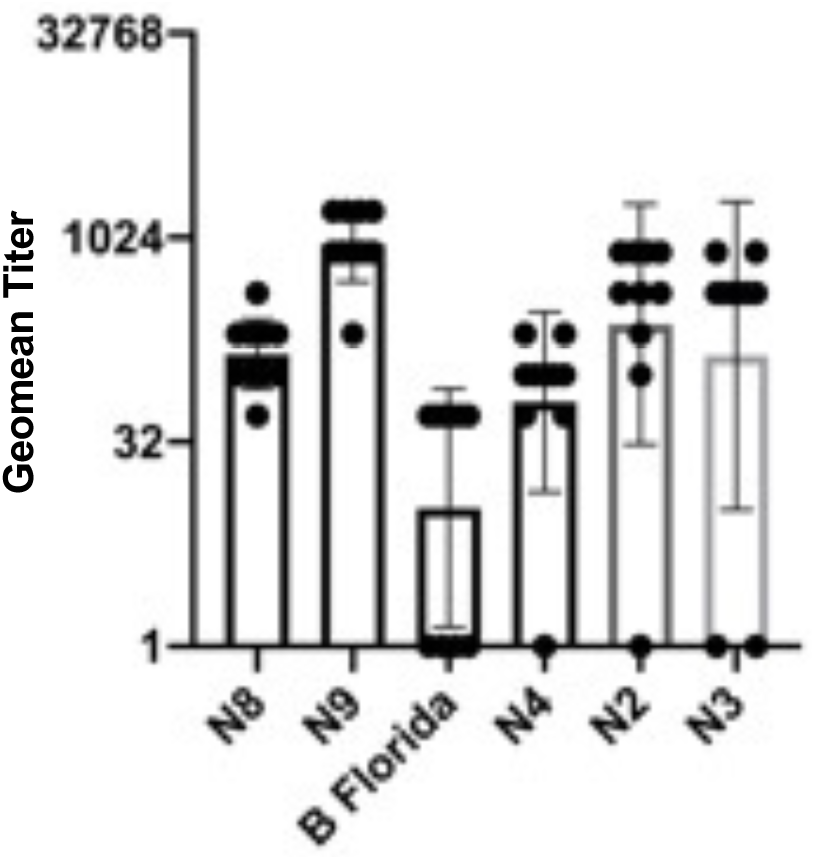

